# Fermented polyherbal formulation ameliorates the severity of acute multiple - antibiotics - resistant *Pseudomonas aeruginosa* infected burn wound in rat burn model

**DOI:** 10.1101/2023.11.13.566953

**Authors:** Subhanil Chakraborty, Subhajit Sen, Ranadhir Chakraborty

## Abstract

*Pseudomonas aeruginosa*, a Gram-negative opportunistic bacterium has emerged as a cause of life-threatening infections in topical burn wounds. Current therapeutic approaches thru wound dressings and systemic medicines are far from satisfactory; multiple-antibiotic-resistance shown by pathogens contribute to failures of therapy causing mortality. This animal study was conducted to check efficacy of one Ayurveda based fermented polyherbal preparation (AP 01) against multiple antibiotic resistant *Pseudomonas aeruginosa HW01* infected rat burn wounds. AP-01 was applied on artificially inflected burn wound on rat model infected with Multiple Antibiotics Resistant *Pseudomonas aeruginosa* to register the healing effects in terms of reduction in residual wound area percentage, presence of C-reactive protein in blood, presence of viable bacteria colony while keeping conventional antibiotics as positive control. The polyherbal preparation had reduced the infected residual burn wound area at 40.63% ± 0.69 from the initial burn wound area within two weeks after a single intervention; whereas residual burn wound area remained much higher in case of animals left untreated and in case of the animals treated with control drug. Restoration to normalcy of serum C-reactive protein level were also achieved earlier in case of polyherbal AP-01 treated groups than other groups. Fermented formulation using components of AP-01 singly or in different combinations were never been tested earlier for topical application in infected burn-wound. The formulation AP-01 was found superior in terms of rate of healing and control of infection by multiple-antibiotic-resistant *P. aeruginosa* strains in burn wounds in rat model.

## Introduction

*Pseudomonas aeruginosa* is a monoflagellated motile, rod-shaped, non-spore forming Gram negative bacterium which has an opportunistic pathogenic nature that causes it to produce serious infections in plants, animals and human beings [1]. People suffering from severe burn injuries as well as patients with other wide varieties of pathogenic states are suffering from Pseudomonas infections in all around globe. Notably inherent broad spectrum antimicrobial resistance in *Pseudomonas aeruginosa* isolates have limited the scope of current antibiotics regimen in treating Pseudomonas infections and as identified by the Centers for Disease Control and Prevention (CDC), USA; as much as 8% of all healthcare-associated infections were found to be caused by *Pseudomonas aeruginosa* isolates, among which about 13% are caused by multidrug resistant strains attributing more than 400 deaths directly per year in USA alone [2]. Among the eight categories of antibiotics normally used in clinical practice to treat *P. aeruginosa* infections, multidrug resistant or multiple antibiotic resistant *Pseudomonas aeruginosa* isolates were identified if it is resistant to more than one therapeutic agent belonging to at least three categories [3].

Ayurveda, the doctrine of ancient Sanskrit scripted resolutions, addressing the health and well-being of human populations and animals had always offered holistic, nature based curative means in treating wide varieties of diseases, disorders and symptoms of medical and pathological importance. Concocting combinations of extracts from several plant parts according to Ayurveda formulary to yield a synergistic therapeutic effect against an illness was a common practice among ancient physicians and healers of India. Possible benefits or threats of the Ayurveda based monoherbal therapy or polyherbal therapy in treating several disorders were tested through pre-clinical and clinical research in different types of diseases with very little scientific proceedings towards enriching the potential of therapeutic benefits into designing modern drugs supported by thorough in-vitro and in-vivo studies [4]. “Arishtha” preparations are particularly important as these are herbal decoctions made with boiling water where certain used plants are capable of self-fermenting and finally produce stable liquors with intended therapeutic property [5]. In the current study, we have demonstrated the potential of Ayurveda based polyherbal preparation (named as AP-01; prepared in the laboratory following the Arishtha manufacturing protocols from Ayurveda Pharmacopoeia) in treating burn wound infections in a rat burn model. The components of AP-01 singly or in different combination of two or all mixed together has never been tested earlier or treated topically to control burn-wound infection what-so-ever. The scientific cue to design experiments to control bacteria-infected-burn-wound arose from the observations that AP-01 was not only able to inhibit swarming and biofilm formation of *Pseudomonas aeruginosa* but also effective in killing multiple-antibiotic-resistant *Pseudomonas aeruginosa* at MIC dose.

Ubiquitous presence of *Pseudomonas aeruginosa* in water, soil and different surfaces has helped it in easily infecting burn wound surfaces and further invasion in systemic circulation [6]. Colonization and persistence on wound surface and resistance towards empirical antibiotics therapy have made it emerge as one of the most threatening infection-causing pathogens in people suffering with burn wounds [7]. Multidrug resistant *Pseudomonas aeruginosa* pathogens are placed in top priority list of World Health Organization posing critical threat to cause morbidity and mortality and also considered under serious threat level by CDC due to the fact that current arsenal of healthcare professionals in treating Pseudomonas infections are falling short day by day [8]. In Europe, 33.9% of *P. aeruginosa* was found to be resistant to at least one of the antimicrobial groups (piperacillin ± tazobactam, fluoroquinolones, ceftazidime, aminoglycosides and carbapenems) under surveillance, according to the report published in 2016 by eCDC or European CDC.^3^ This study highlights on findings related to treat MAR *Pseudomonas aeruginosa* infected burn wound and further control of infection spreading through systemic circulation with the help of fermented polyherbal Ayurveda based formulation – AP-01.

## Materials and methods

### Crude plant parts for the fermented polyherbal formulation

Fermented polyherbal formulation (AP-01) was prepared in laboratory following Ayurveda; by boiling dried and cleaned stem barks of *Holarrhena antidysenterica* & *Gmelina arborea* mixed with large sized dry resins of *Vitis vinifera* in presence of dried flowers of *Madhuca indica* in potable drinking water. Dried flowers of *Woodfordia fruticosa* [9] was added after the volume of water lowered to one quarter of the initial volume thru boiling for initiation of fermentation while using Jaggery as a source of sugar [5]. After fermentation completed in about 45 days, the formulation was duly filtered and kept in sterile dark glass bottles after passing through bacterial filters of 0.2-micron pore size for further use. The method of preparation was in accordance with The Ayurvedic Pharmacopoeia of India following references of Sharangadhar Samhita and Bhaisajyaratnavali for different “Arishta” formulations [10–12]. Antibacterial activity was tested for each active plant parts of *Holarrhena antidysenterica*, *Gmelina arborea*, *Vitis vinifera* and *Madhuca indica* for equivalent strength by using *Woodfordia fruticosa* for fermentation and Jaggery as a source of sugar to make arishtha preparations from individual components of the polyherbal preparation. All plant parts were procured from local market and identified by qualified Ayurveda practitioner bearing formal academic degree for practicing Ayurveda. Safety profile of the polyherbal Arishtha preparation was established from the literature and the physician for both oral and topical routes of administration.

Piperacillin/tazobactam (Lupin ltd., India) was used as a standard drug in animal study related to infected burn wound healing by polyherbal preparation AP-01.

### Experimental animals

Thirty-six (36) Sprague Dawley rats of either sex and average six weeks old, weighing 120 to 160gms were obtained from animal house of Department of Zoology, University of North Bengal and distributed randomly into six groups where every group contained equal numbers of male and female animals. Animals were acclimatized for seven days in standard environmental condition and pellet diet of chow and water was provided ad libitum.

### Microorganism

The strain *Pseudomonas aeruginosa* HW01 isolated from the district hospital waste water [13] was used to induce external infection on burn wound site in the study of *in-vivo* wound healing assay.

### Antimicrobial Susceptibility Test

Antibiotic resistance/susceptibility profile of *P. aeruginosa HW01* was determined as per the guidelines of CLSI (National Committee for Clinical Laboratory Standards Institute) [14]. Kirby-Bauer disk-diffusion method was applied to determine the antibiotic resistance/susceptibility profile of *P. aeruginosa HW01*. Mid log-phase culture of HW01 (equivalent to 0.4 McFarland standard) was spread on Muellar-Hinton agar (MHA) plates by spread-plate technique. After spreading the culture evenly over the entire surface of the media, different antibiotic disks (HiMedia, India) such as, amikacin-AK (30 μg), aztreonam-AT (30 μg), cefepime-CPM (30 μg), ceftazidime-CAZ (10 μg), ciprofloxacin-CIP (5 μg), colistin-CL(10 μg), doripenem-DOR (10 μg), levofloxacin-LE (5 μg), meropenam-MRP (10 μg), piperacillin/tazobactam-PIT (100/10 μg), ticarcillin-TI (75 μg), and tobramicin-TOB (10 μg) were gently placed on the plates and incubated for 24 h at 37°C. After incubation, the zone of inhibition was measured and compared with break-points given by EUCAST/NCCLS to determine the resistance/susceptibility profile of *P. aeruginosa HW01* against above mentioned antibiotics.

### Antimicrobial Assay of polyherbal AP01 against P. aeruginosa HW01

The minimum inhibitory concentration (MIC) of polyherbal preparation AP01 against the *P. aeruginosa HW01* was determined by broth dilution method followed by National Committee for Clinical Laboratory Standards Institute (CLSI) [14]. Polyherbal preparation AP01 was added to sterile 5ml Muller Hinton broth in different concentration from 0 µl/ml to 150 µl/ml with a gradual increment of 10 µl/ml and then 1% aliquot of mid log-phase culture of *P. aeruginosa HW01* was inoculated in each tube equally and incubated for 24 h at 37°C. After incubation, growth was measured in spectrophotometer (SPECTROSTAR Nano - BMG LABTECH) at 600nm to determine the minimum inhibitory concentration (MIC). All the experiments were done in triplicates and repeated twice. Overnight grown mid-log phase *P. aeruginosa* culture was inoculated both in presence of polyherbal AP-01 at MIC dose and without the presence of polyherbal AP-01 at 37°C and incubated for six hours to compare the optical density-based growth in both conditions using the spectrophotometer.

### Swarming motility assay of Pseudomonas aeruginosa HW01

Swarming motility assay of *P. aeruginosa HW01* was carried out on soft agar medium following standard method [15]. Luria Bertani (LB) broth was supplemented with 0.5% agar and 5.0 % glucose to prepare soft agar medium. Liquid polyherbal preparation AP-01 at a volume of 40µl per ml media was added to sterile soft agar medium in prior to pouring the test plates. In the case of preparing control plates, sterile distilled water (equivalent to the volume of AP01 added in the test medium) was added to sterile soft agar medium before pouring the plates. 10 µl of the overnight grown culture of *P. aeruginosa HW01* was inoculated in the centre of the test and control soft agar plates and incubated at 37°C for at least 18 h in the upright position to compare the diameter of the swarming zone in both test and control plates.

### In-vivo wound healing assay

*In-vivo* wound healing experiment was done by following our previously published research work, in accordance with National Research Council’s Guide for the care and use of laboratory animals [16–17]. The study was approved by the Institutional Animal Ethics Committee of University of North Bengal, Siliguri, Distr. Darjeeling, West Bengal, India (Ref. No. IAEC/NBU/2018/04 dated 12.09.2018). Parallel animal studies were done with six Sprague Dawley rats randomly assigned into each group, maintaining the sex ratio of 1: 1, and one animal was kept per vinyl cage (Tarson, India) to prevent possible aggression, if any, of one animal towards the other of the same sex or opposite. The bottom of the cages was lined with rice husk and were kept in animal enclosure under controlled atmospheric conditions of temperature (25 +/− 2 °C), humidity (55 +/− 5%), and a diurnal variation of 12 hours light and dark period were maintained throughout the study period. Food pellets of standard chow diet (Pranav Agro Pvt.Ltd., India) and machine filtered tap water (Aquaguard, Eureka Forbes, India) were fed to the animals ad-libitum.

### Burning procedure

Slight modifications were applied on the method of burning techniques reported by earlier authors to develop an animal burn model [18–19]. Site for burning was chosen at dorsal side of rats while keeping equivalent proximity from heads and other internal vital organs. The furs of the rats were shaved from the site on back, covering equal area from spinal cord at first with the help of sterile scissors and razor. General anesthesia was performed using a combination of ether chloroform at a ratio of 1:3 v/v in prior to placing a piece of fire-proof clothing, carefully over the site chosen for inflicting burn. The site of intended burn was kept free and visible through a cautiously cut window (at the fire-proof clothing) of one square inch area, which was poured with 0.2 ml of 95% ethanol, while the rest portion of the clothing was held tightly against the rat body surfaces. The poured ethanol was lit up and left to burn and the flame was extinguished exactly after 15 seconds. Intra-peritoneal injection of 0.5 ml physiological saline solution (PSS; 0.85% NaCl) was injected to each animal immediately after the burn episode to revive them from the shock of burn and anesthesia while they gain consciousness.

### Topical infection on the burn-wound site

Animals belonging to three groups were infected by inoculating the wound site by *Pseudomonas aeruginosa HW01* hospital waste water isolate [13]. At first, cell pellets were obtained by centrifuging overnight grown *P. aeruginosa HW01* cells in Luria broth (Himedia, India) at 4000 revolutions per minute at 40 Centigrade temperature and then the cell pellet was resuspended in sterile phosphate buffer saline (PBS) after two consecutive washes with PBS to obtain a cell density of 109 bacterial cells per milliliters.

An aliquot of 200 μl cell suspension from the same tube was used to infect single burn zone in each animal. A single layer of sterile non-absorbent cotton gauze was temporarily placed over the burn-area for 10 minutes to aid effective absorption of the inoculums before application of any other treatment [16]. The weights of the rats were taken on a daily basis and the food intakes by rats were also noted regularly.

### Topical treatment of burn-afflicted site

All treatments were provided after infecting the wound sites with *P. aeruginosa HW01* and removal of the cotton gauge. Animals were already divided into six groups, each group comprising six animals where a male to female ratio of 1:1 was maintained. Animals of group I, II and III were all inflicted with burn but the burn site was not interfered with *P. aeruginosa HW01* infections, whereas all animals from group IV, V and VI were bearing *P. aeruginosa HW01* infections over their burn wounds. Burn wound of group I was given 100 µl distilled water. Burn wound of group II was treated with 100 µl of piperacillin/tazobactam solution where each ml contained 56/7 µg/ml piperacillin/tazobactum as *P. aeruginosa HW01* was found susceptible to this combination. Group III animals were treated with 100 µl of AP-01 spread over the wound surface. Infected burn wound areas of group IV animals were spread with 100 µl distilled water and the infected burn wound sites of group V and group VI animals were respectively treated with piperacillin/tazobactam and polyherbal AP-01. All the thirty-six rats, distributed in six different groups undergoing six different treatments were observed for two weeks on a daily basis till disappearance of C-reactive proteins and count of viable Pseudomonas colonies from infected wound site (explained in later part of this paper) and for those two weeks, percentages of residual wound area were physically measured to reciprocate possible wound healings quantitatively for statistical validation [16].

### Qualitative determination of the pathogen load in the burn-wound site

Microbiological evaluation was undertaken from the next day of the burning experiment by using “sterile swabs” from the site of infected burn wounds and burn wounds without addition of *P. aeruginosa* inoculums as well. The burn wound exudates were collected with the cotton swabs in prior to dipping the entire swab in 1ml sterile PBS solution. 100 µl aliquot from the swab drained PBS was then aseptically spread over King’s Medium A Base (HIMEDIA, M1543-500G) plate. Spread-plates in triplicates were evaluated for *Pseudomonas aeruginosa HW01* colonies after 24 h of incubation. This evaluation was continued routinely until the disappearance of *Pseudomonas aeruginosa HW01* colonies occurred from infected wound sites [16].

### Determination of C-reactive protein (CRP) levels post burn and infection

Certain volumes of blood samples (15 to 20 µl) of the rats were collected in prior to intervention and after every 48 hours intervals from the tail veins, saphenous veins and dorsal pedal veins in the sterile 0.25 mL PCR tubes for determination of underlying infections present in the animals. Blood sample containing tubes were placed vertically for a period of 30-45 mins for settling down the blood corpuscles. The blood samples were then centrifuged for 5 min at 3600 rpm in order to separate the serum. Sterile PCR tubes were used for collecting the supernatant serum samples from the blood. The collected serum samples were used to determine qualitative and semi-quantitative C-reactive protein (CRP) levels using RHELAX-CRP latex reagent (Tulip Diagnostics (P) Ltd., India). Physiological saline solution was prepared for preparing the testable dilutions of the serum from different specimens. 1:2, 1:4, 1:8, 1:16, 1:32 dilutions were made for quantitative CRP measurement and RHELAX-CRP latex reagent was mixed with diluted samples as per the directions provided in instructions manual. Then the mixture was mixed properly by the mixing stick and the agglutination reaction was observed macroscopically within 2-3 min for proper determination of CRP levels of each animal subjected to experiment [16].

### Statistical analysis

Thirty-six burn-injured animals were equally divided into six groups (3 male + 3 female rats/group), Gr-A (burn treated with sterile distilled water alone), Gr-B (burn infected with *P. aeruginosa HW01*), Gr-C (burn treated with piperacillin/ tazobactam), Gr-D (burn infected with *P. aeruginosa HW01* and treated with piperacillin/ tazobactam), Gr-E (burn treated with AP01), and Gr-F (burn infected with *P. aeruginosa HW01* and treated with AP01). Post-treated residual burns wound area, expressed as percent residual wound areas (PRWA) per treatment group on different day following treatment (0/2nd/4th/6th/8th/10th/12th/14th day) were used to determine mean and standard deviation. The p-value corresponding to the F-statistic of one-way ANOVA was determined in order to suggest whether 14th day data on PRWA from one or more treatments were significantly different. This was followed by Tukey HSD test, Scheffe, Bonferroni and Holm multiple comparison tests (astatsa.com/OneWay_Anova_with_Tukey HSD/). These post-hoc tests were intended to identify pairs of treatment that are significantly different. immediate release microspheres.

## Results

### Antimicrobial susceptibility test

As per the antimicrobial susceptibility tests conducted following CLSI guidelines, the hospital wastewater isolate *Pseudomonas aeruginosa HW01* was found resistant to ten out of eleven tested antibiotics from different classes of antibiotics (Table 1). It was also found, that the isolate was susceptible to piperacillin/tazobactam combination marginally.

**Table 1.**
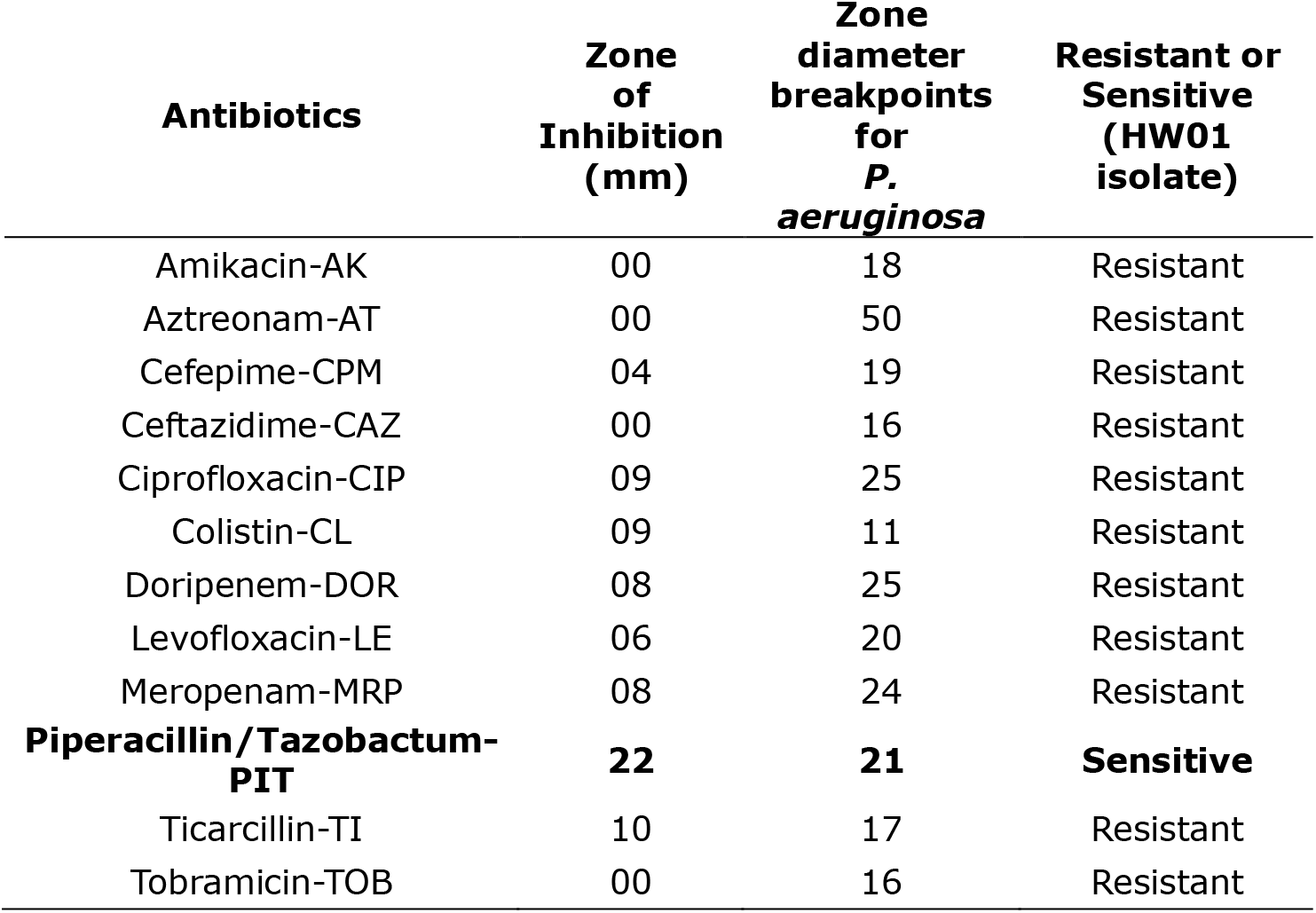
Antibiotic resistance profile of *Pseudomonas aeruginosa HW01*.

### Swarming motility assay of Pseudomonas aeruginosa HW01

*P. aeruginosa HW01* formed dendrite like outward projections from the center point of bacterial inoculation over semi solid media as the branching occurred due to the swarming motility of the bacteria (Fig. 1A). In presence of sub-MIC dose of 40µl per ml polyherbal AP-01 in semi-solid media, the Fig. 1B showed that the bacterial cells were able to grow and form a disoriented colony at the center but were unable to produce extensive branching or tendrils suggestive of inhibition of swarming motility due to the action of AP-01.

**Fig. 1.**
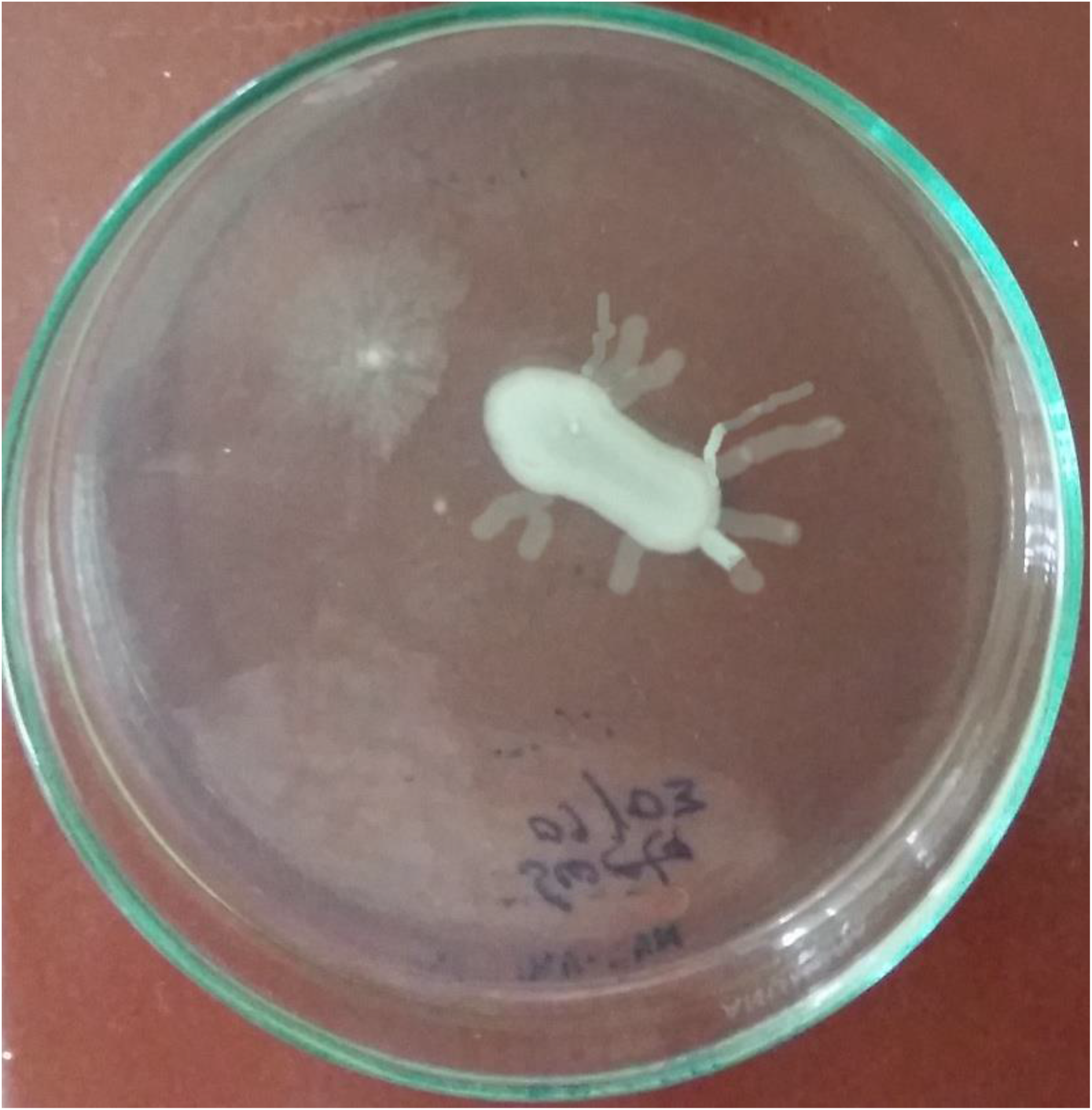
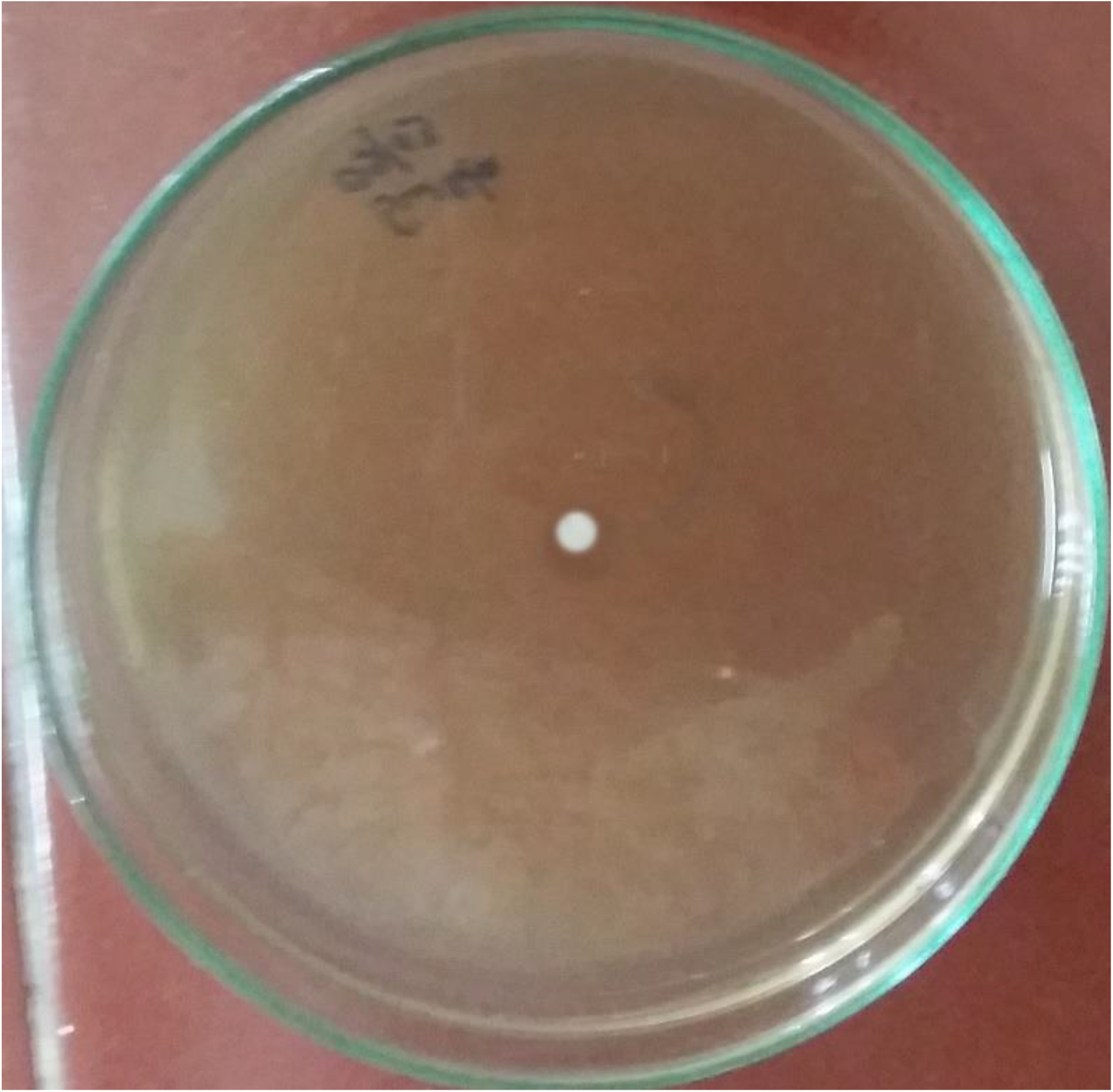
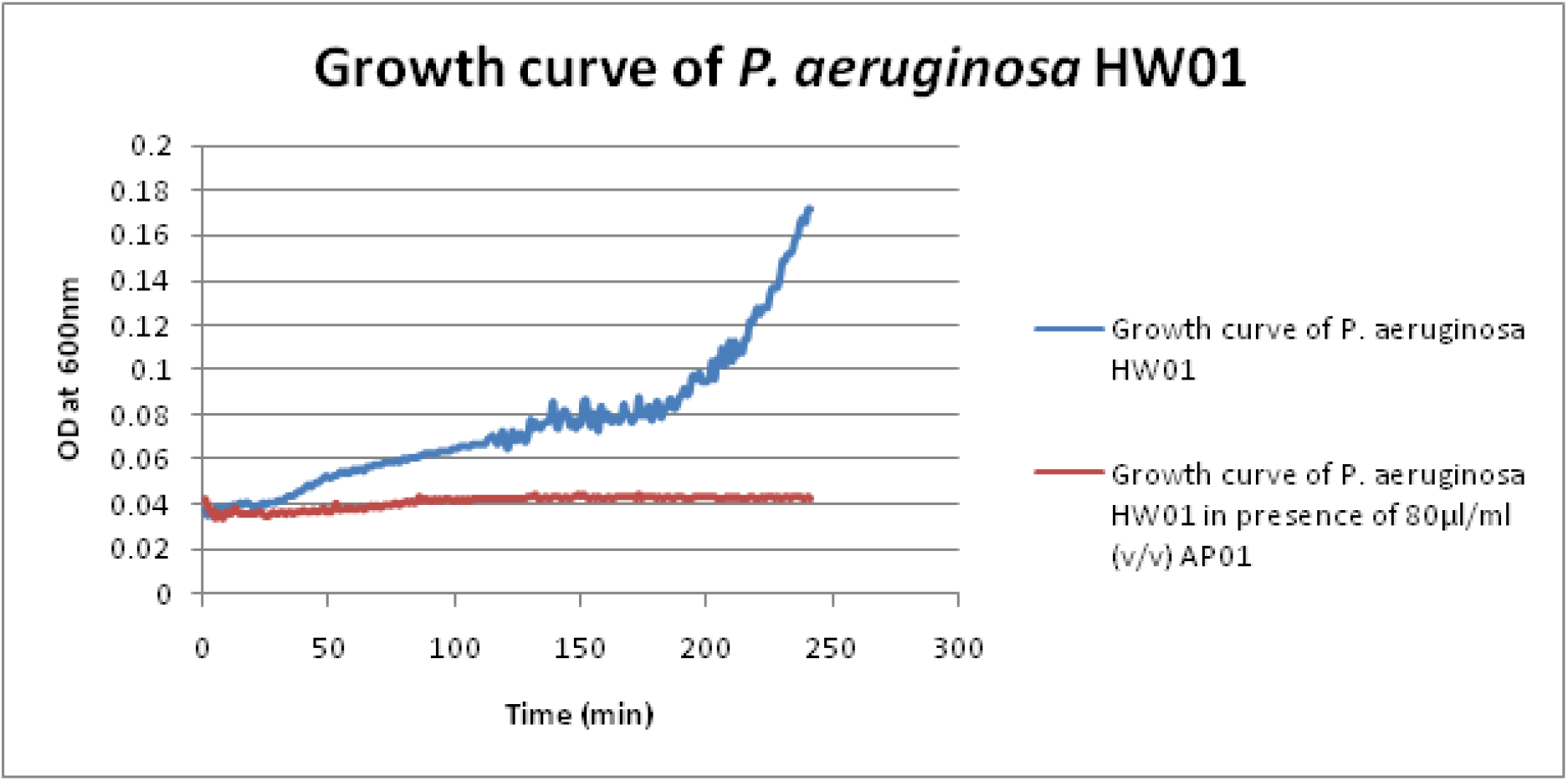
(Color) Swarming motility assay of *P. aeruginosa HW01*: **(A)** Control (in absence of AP-01), **(B)** in presence of polyherbal preparation AP-01 and **(C)** Growth curve of *P. aeruginosa HW01*, in presence of 80 µl/ml (v/v) polyherbal preparation AP-01 and in absence of AP-01.

### Antimicrobial Assay of polyherbal AP-01 against P. aeruginosa HW01

No visible growth was observed through spectrophotometric method after 24 hours incubation at 37°C from the tube containing 82 µl per ml AP-01 and higher drug volumes onwards (Data not shown). Therefore, the minimum inhibitory concentration of polyherbal AP-01 was determined to be 82 µl per ml against *Pseudomonas aeruginosa HW01*. Individual components of the polyherbal formulation were assayed separately to check the potentials of the antibacterial effect against *Pseudomonas aeruginosa HW01* as shown in Table 2 suggested better efficacy in terms of MIC shown by polyherbal formulation AP-01 than other individual components. Fig. 1C depicted no visible growth of *Pseudomonas aeruginosa HW01*, when it was grown in presence of minimum inhibitory concentration of AP-01 in Mueller Hinton Broth for the first six hours after inoculation with mid-exponential phase cells. However, in absence of AP-01, the bacterial cells were found to be growing steadily as shown in the same figure (Fig. 1C).

**Table 2.**
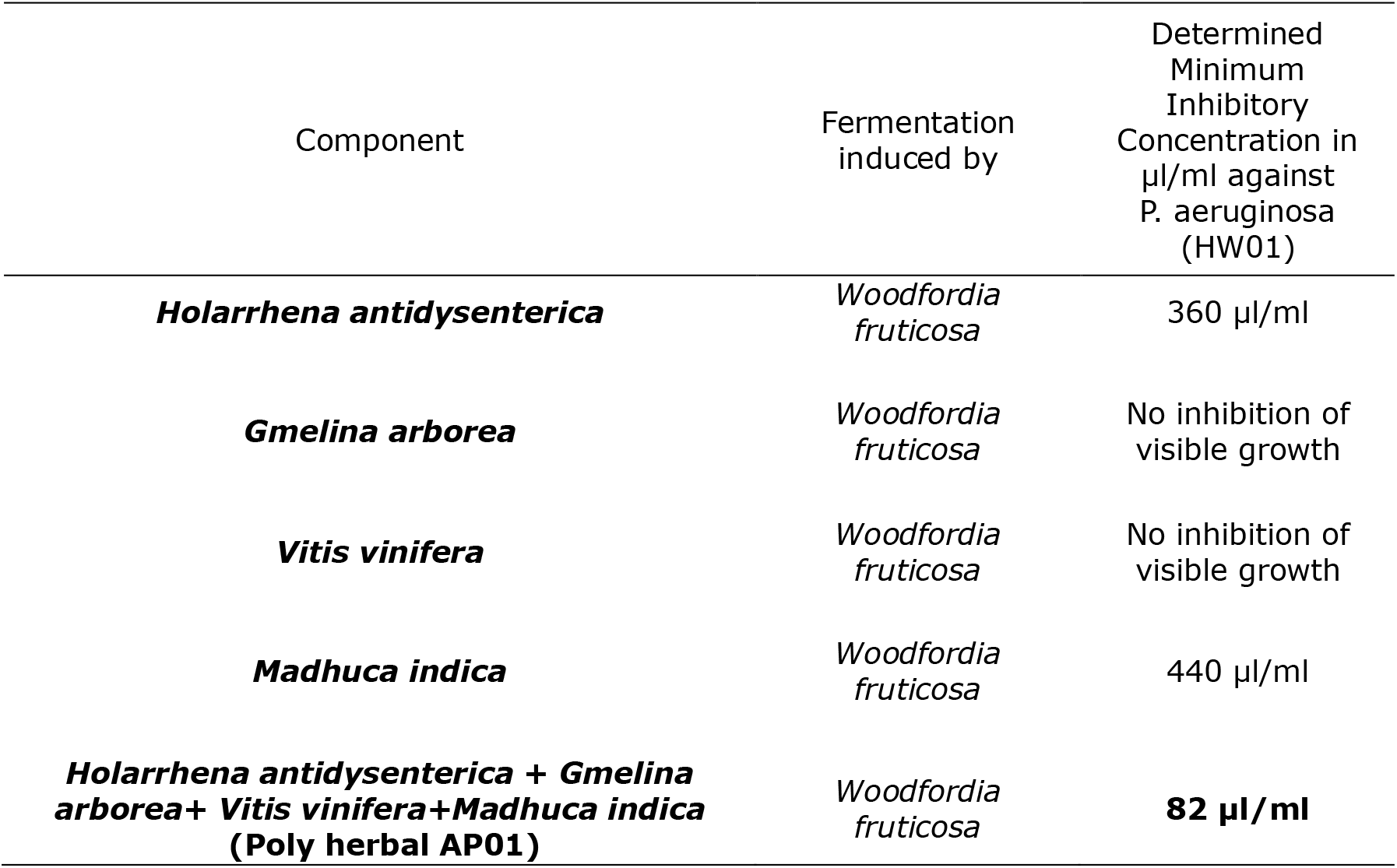
Comparative analysis and determined MIC of individual component and the polyherbal AP-01 against *Pseudomonas aeruginosa HW01*.

| Component | Fermentation induced by | Determined Minimum Inhibitory Concentration in µl/ml against *P. aeruginosa* (HW01) |
| --- | --- | --- |
| ***Holarrhena antidysenterica*** | *Woodfordia fruticosa* | 360 µl/ml |
| ***Gmelina arborea*** | *Woodfordia fruticosa* | No inhibition of visible growth |
| ***Vitis vinifera*** | *Woodfordia fruticosa* | No inhibition of visible growth |
| ***Madhuca indica*** | *Woodfordia fruticosa* | 440 µl/ml |
| ***Holarrhena antidysenterica* + *Gmelina arborea* + *Vitis vinifera* + *Madhuca indica* (Poly herbal AP01)** | *Woodfordia fruticosa* | **82 µl/ml** |

### In-vivo wound healing assay

Greater re-epithelization and reduction of wound area and reappearance of furs in the burn wound area were also evident in case of animals treated with polyherbal AP-01 in both infected and non-infected burn groups (Fig. 2A & 2B).

**Fig. 2.**
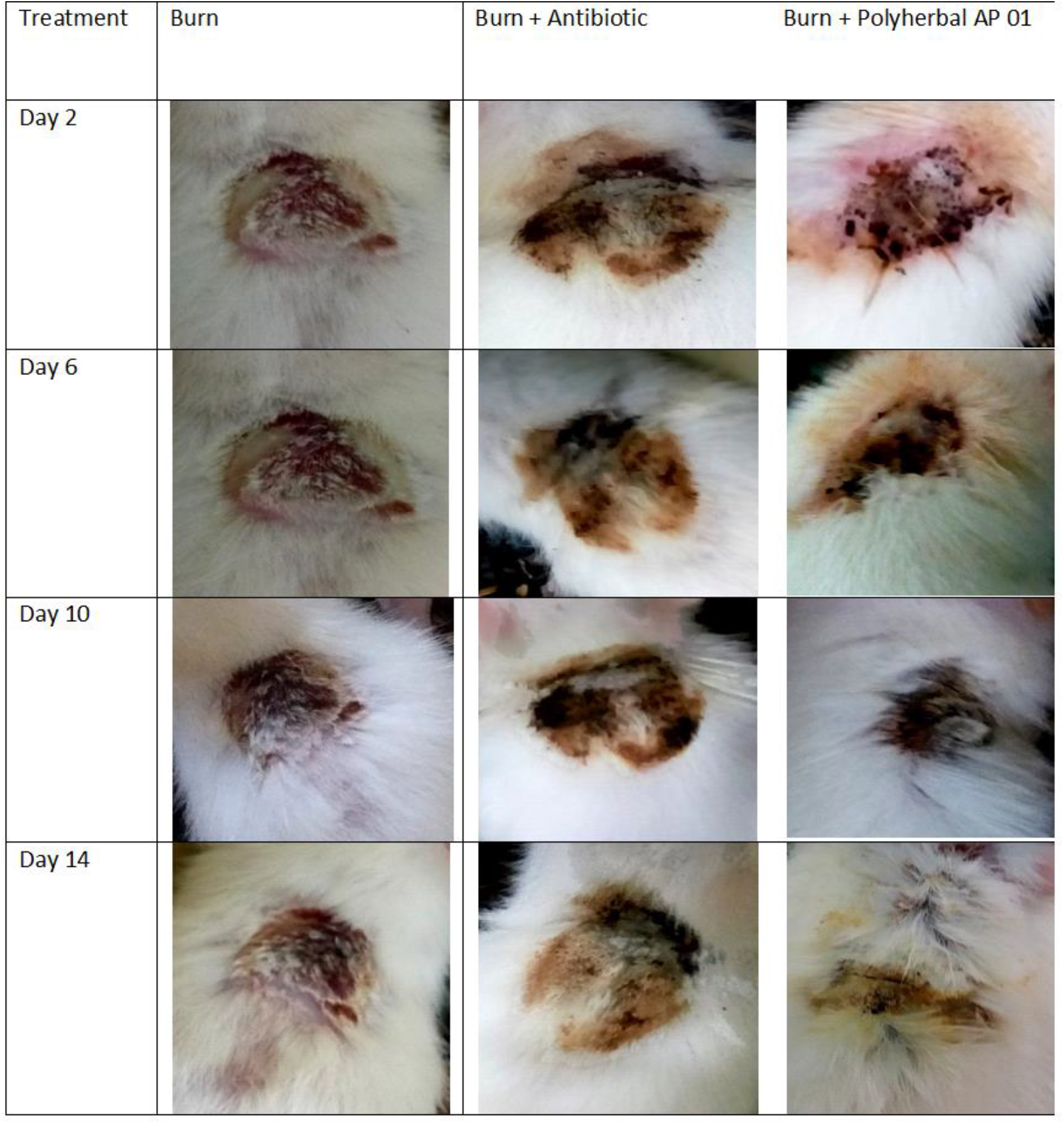
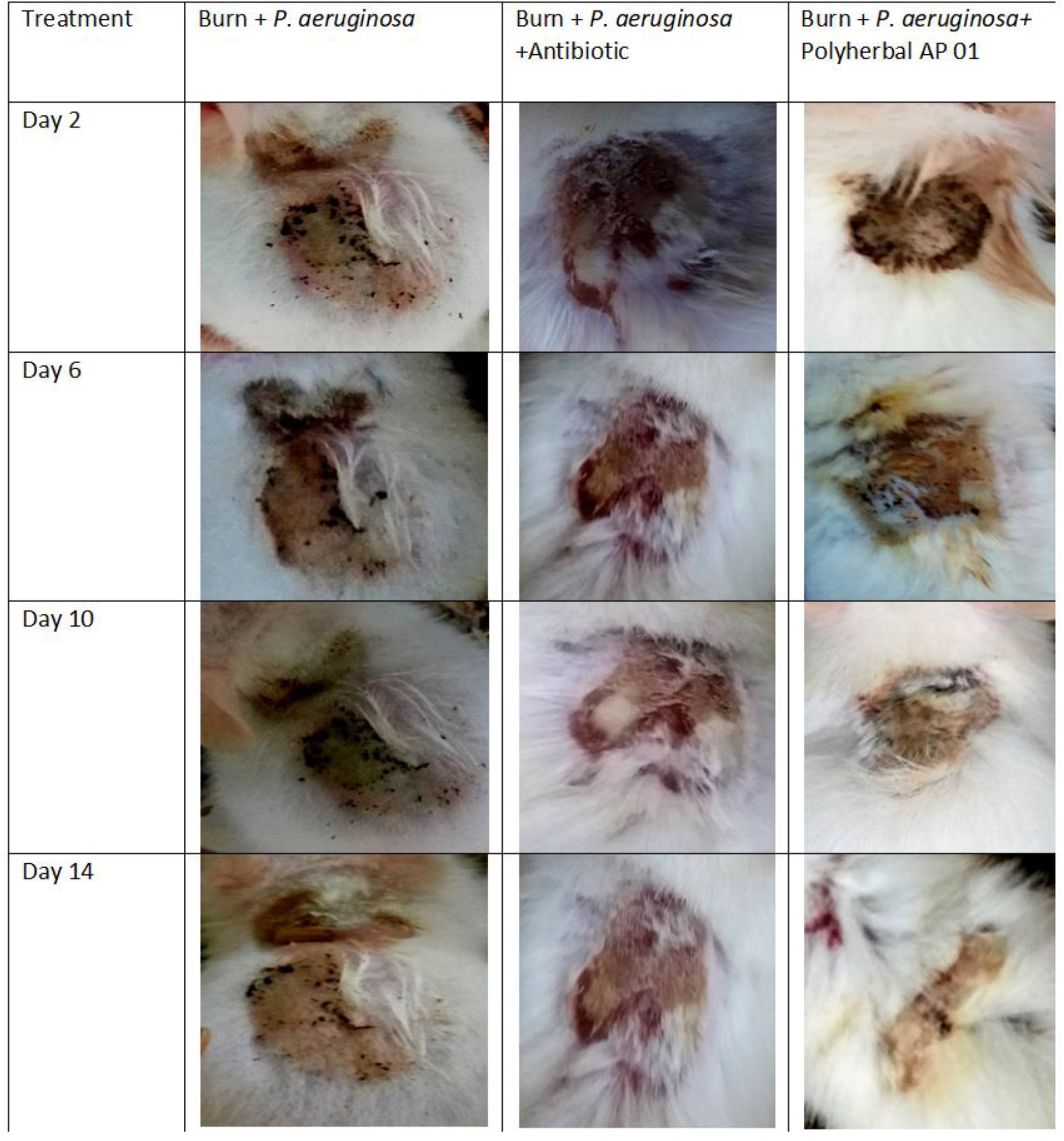
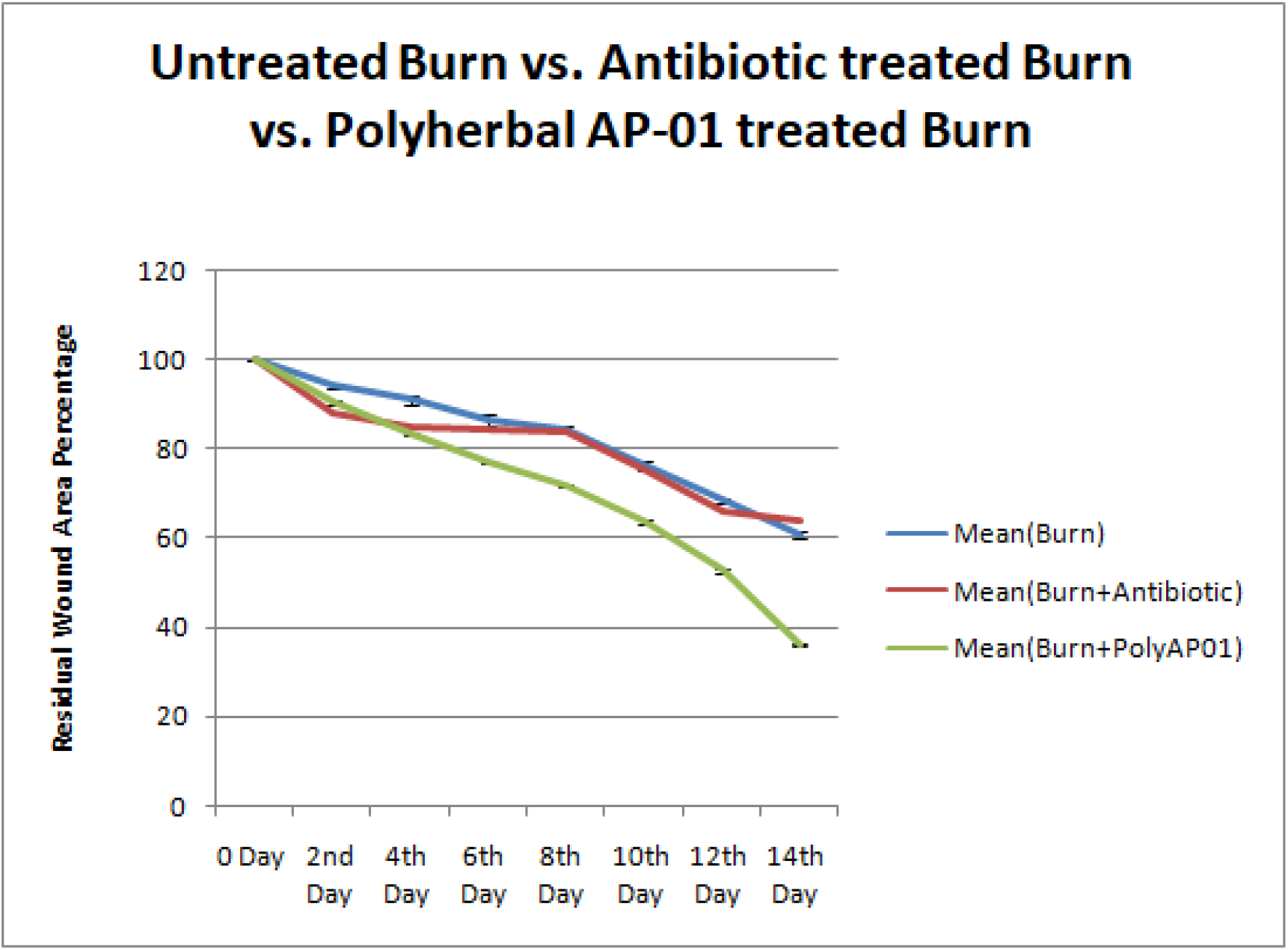
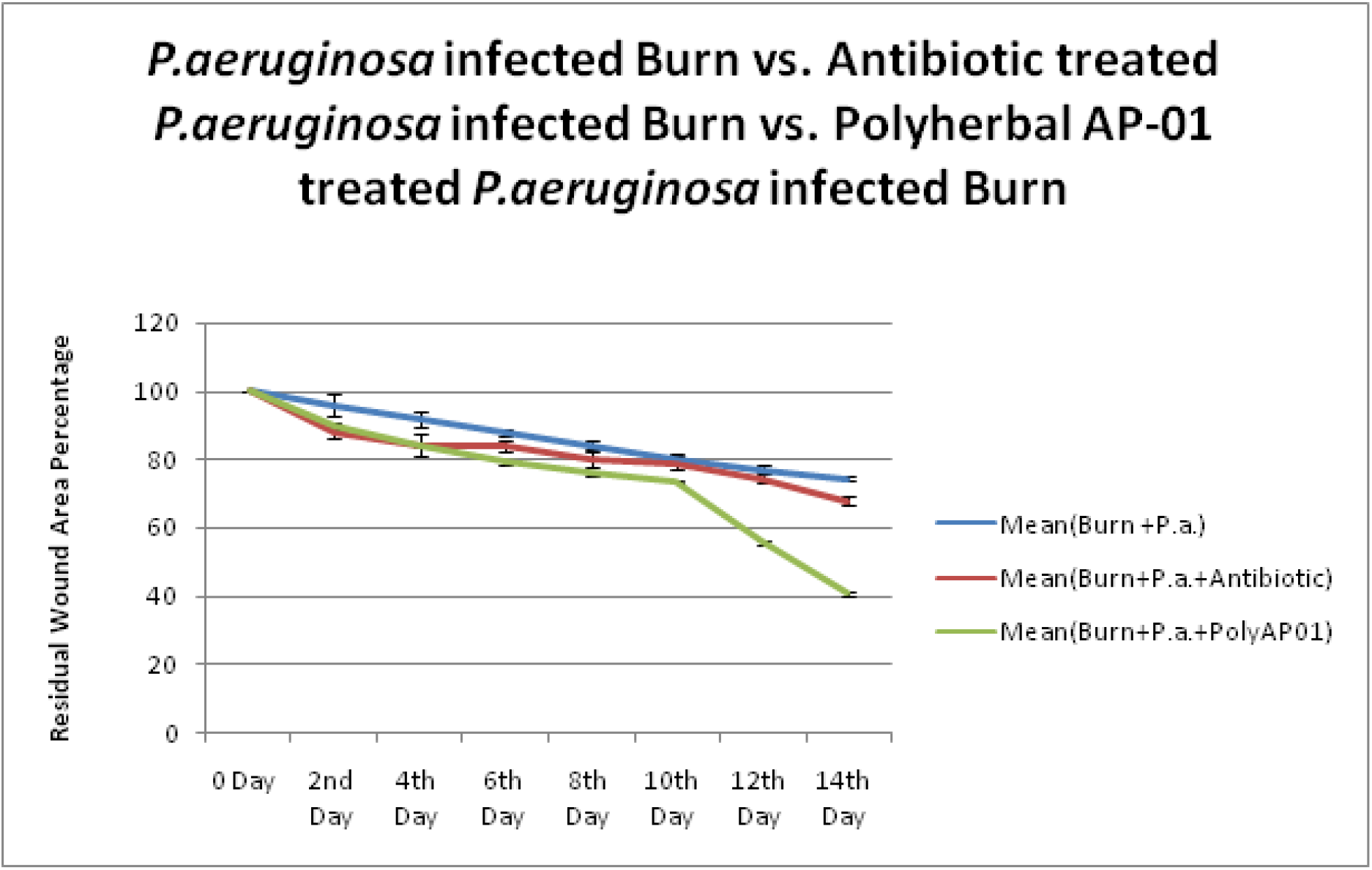
(Color) Decrease in residual wound area with progression of burn wound healing: **(A)** untreated burn (first column), burn treated with antibiotics (second column), burn treated with polyherbal preparation AP01(third column), **(B)** *P. aeruginosa HW01* infected burn (first column), *P. aeruginosa HW01* infected burn treated with antibiotic (second column), *P. aeruginosa HW01* infected burn treated with polyherbal preparation AP-01(third column) and graphical representation of in vivo wound healing assay, **(C)** comparison of residual wound area from day 0 to day 14 among Untreated burn wound, antibiotics treated burn wound and polyherbal AP-01 treated burn wound. **(D)** comparison of residual wound area from day 0 to day 14 among untreated infected burn wound, antibiotics treated infected burn wound and polyherbal AP-01 treated infected burn wound.

An incomplete consistency over the burn wound surface area (considered 100% for each rat) was attained from flame burning technique applied in the current study. Animals from groups I, II and III, where the burn wound was not infected with *P. aeruginosa HW01*; the decrease in residual wound area was found to be maximum in group III animals receiving AP-01 treatment and it was determined to be 36.12% ± 0.46 (Fig. 2C). Animals from groups IV, V and VI where the burn wound was infected with *P. aeruginosa HW01*, the decrease in residual wound area was also found to be maximum in group VI animals receiving AP-01 treatment and it was determined to be 40.63% ± 0.69 (Fig. 2D).

It was observed that the burn injury with simultaneous infection of *P. aeruginosa HW01* caused log-fold increase from the normal level of serum C-reactive protein on day 2 after burn injury in both treated and untreated animals irrespective of intervention with *P. aeruginosa* infection. The increased CRP level was receded faster to normal level with control of inflammation in the animals treated with polyherbal AP-01 (group III and group VI) within day 8 when serum CRP became undetectable. Whereas, increased level of CRP could be evidently detected in animals of other groups (group I, II, IV and V) as shown in (Table 3).

**Table 3.** Level of C-reactive protein (CRP) in the blood of different group animals on second, fourth, sixth-and eighth-day following burn and infection with *P. aeruginosa HW01*.

| Treatment | Agglutination in different dilutions of the serum |  |  |  |  |  |  |
| --- | --- | --- | --- | --- | --- | --- | --- |
|  | 1:1 | 1:2 | 1:4 | 1:8 | 1:16 | 1:32 | mg/dl |
| 02-day |  |  |  |  |  |  |  |
| Burn | + | + | + | + | + | + | 90-80 |
| Burn + *P. aeruginosa* | + | + | + | + | + | + | 90-80 |
| Burn + Antibiotic | + | + | + | + | + | - | 70-60 |
| Burn + *P. aeruginosa* + Antibiotic | + | + | + | + | + | - | 70-60 |
| Burn + Polyherbal AP-01 | + | + | + | + | - | - | 60-50 |
| Burn + *P. aeruginosa* + Polyherbal AP-01 | + | + | + | - | - | - | 50-40 |
| 04-day |  |  |  |  |  |  |  |
| Burn | + | + | + | + | + | - | 70-60 |
| Burn + *P. aeruginosa* | + | + | + | + | + | - | 80-70 |
| Burn + Antibiotic | + | + | + | + | - | - | 60-50 |
| Burn + *P. aeruginosa* + Antibiotic | + | + | + | + | - | - | 60-50 |
| Burn + Polyherbal AP-01 | + | + | + | - | - | - | 50-40 |
| Burn + *P. aeruginosa* + Polyherbal AP-01 | + | + | - | - | - | - | 40-30 |
| 06-day |  |  |  |  |  |  |  |
| Burn | + | + | + | + | - | - | 60-50 |
| Burn + *P. aeruginosa* | + | + | + | + | + | - | 70-60 |
| Burn + Antibiotic | + | + | + | + | - | - | 60-50 |
| Burn + *P. aeruginosa* + Antibiotic | + | + | + | + | - | - | 60-50 |
| Burn + Polyherbal AP-01 | + | + | - | - | - | - | 40-30 |
| Burn + *P. aeruginosa* + Polyherbal AP-01 | + | + | - | - | - | - | 40-30 |
| 08-day |  |  |  |  |  |  |  |
| Burn | + | + | + | - | - | - | 50-40 |
| Burn + *P. aeruginosa* | + | + | + | + | - | - | 60-50 |
| Burn + Antibiotic | + | + | - | - | - | - | 40-30 |
| Burn + *P. aeruginosa* + Antibiotic | + | + | + | - | - | - | 50-40 |
| Burn + Polyherbal AP-01 | - | - | - | - | - | - | ND* |
| Burn + *P. aeruginosa* + Polyherbal AP-01 | - | - | - | - | - | - | ND* |

\*ND- Not Detectable

The pathogen-load for the particular strain of *P. aeruginosa HW01* gradually reduced to zero number of colonies on the 5th day following burn in case of AP-01 treated animals of group no.VI, while detectable numbers of viable bacteria were still present over the infected burn wounds of either untreated group or the group treated with piperacillin/tazobactam (Table 4).

**Table 4.**
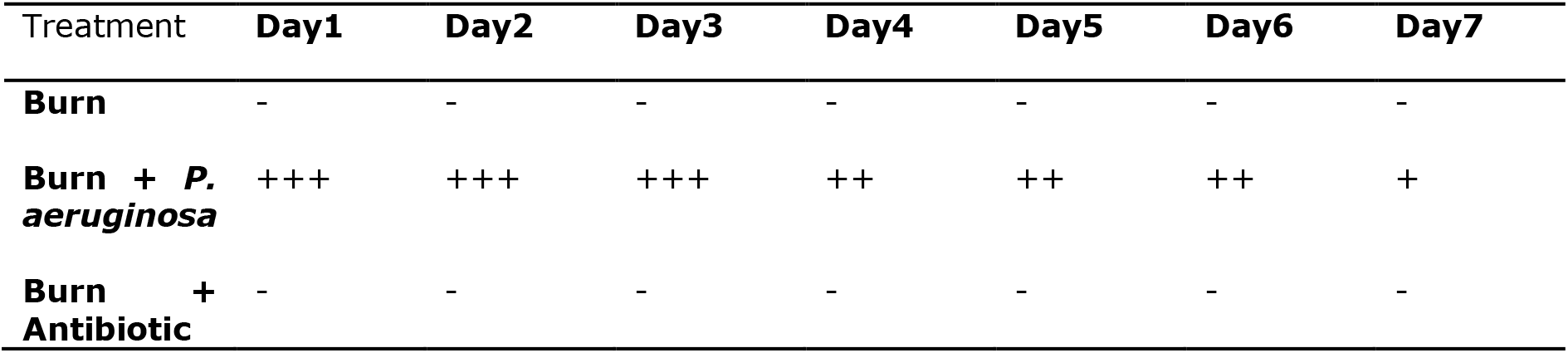
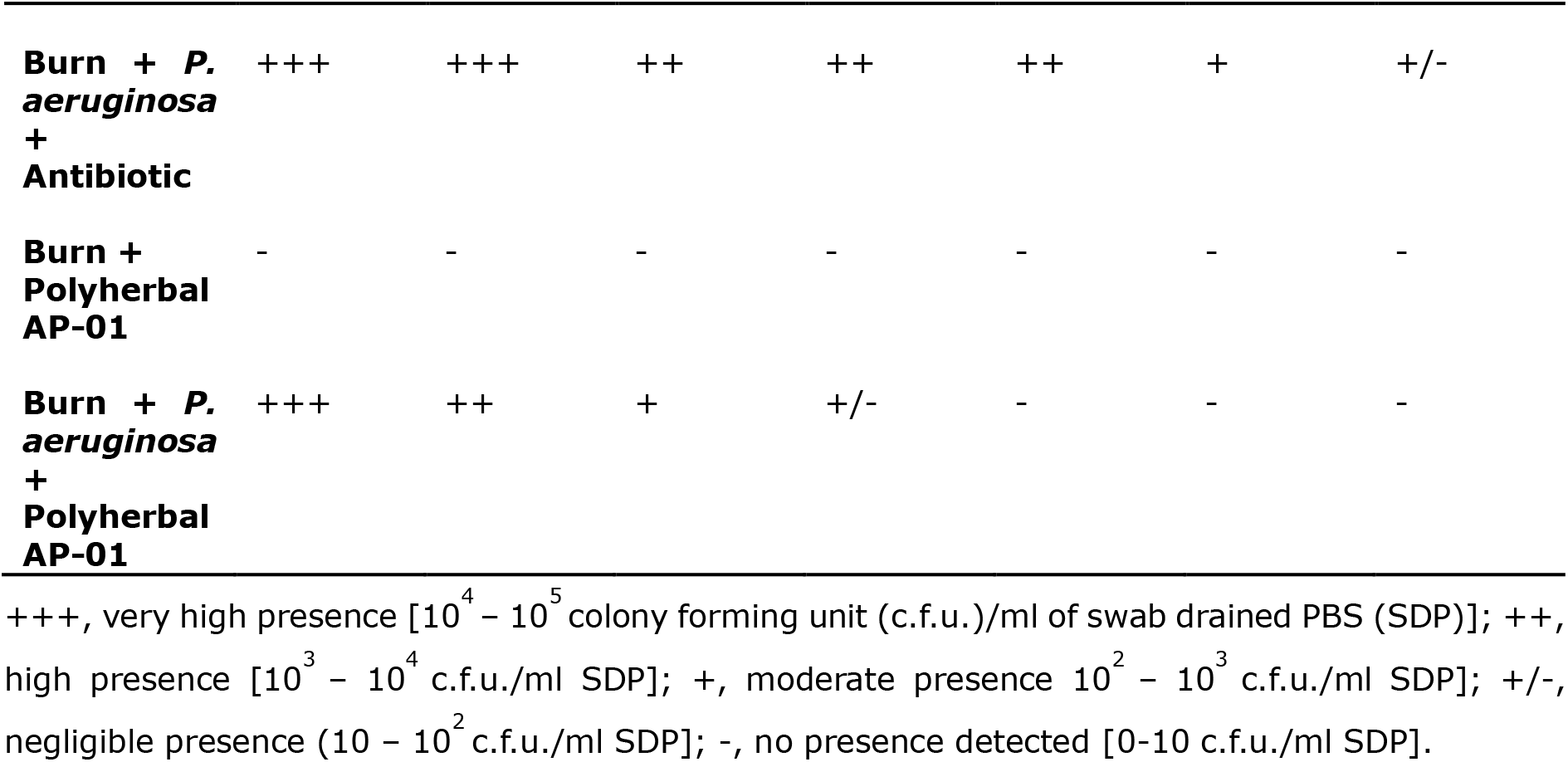
Presence of *P. aeruginosa* viable colonies in seven consecutive days of post-burn wound surface in experimental rats.

### Statistical analysis

The p-value corresponding to the F-statistic of the one-way ANOVA was found lower than 0.01 which strongly suggested that one or more pairs of burn treatments are significantly different. This was followed by establishing a Tukey test statistic from our sample columns to compare with the critical value of the standardized range distribution. Post-hoc Tukey HSD test calculator results have revealed significant difference in the percent residual wound area (PRWA) on the 14th day in the following pairs of treatments. Tukey HSD p-value for the pairs, Gr-I (burn area treated with sterile distilled water) vs Gr-II (burn infected with *P. aeruginosa HW01*) or Gr-II vs Gr-V (burn treated with AP01) or Gr-II vs Gr-VI (burn infected with *P. aeruginosa HW01* and treated with AP01) was 0.0010053. Gr-V vs Gr-VI had least Tukey HSD Q statistic compared to other treatment pairs (mentioned above) yet produced Tukey HSD p-value = 0.0010053. Scheffe multiple comparison results revealed that the Scheffe p-value of the treatment pairs I vs II or I vs V or I vs VI or II vs V or II vs VI was 1.1102e-16 while the pair V vs VI has shown Scheffe p-value = 5.519e-10 (Scheffe inference **p < 0.01). When only pairs relative to I were simultaneously compared, Bonferroni and Holm results have shown that I vs II, I vs V, and I vs VI have yielded highly significant differences with Bonferroni p-value and Holm p-value of 0.000e+00. Such a situation is relevant when treatment A is the control and the experimenter is interested only in differences of treatments relative to control.

Though treatment of burn (with or without *P. aeruginosa HW01* infections) with burn treatment with antibiotics (piperacillin/tazobactam) produced significant differences but Bonferroni p-value and Holm p-value differed widely: I vs II (both Bonferroni p-value and Holm p-value is 0.0000e+00); I vs III (burn treated with antibiotic) yielded Bonferroni p-value = 1.0086e-06 and Holm p-value = 1.6810e-07; I vs IV (burn infected with infected with *P. aeruginosa HW01* and treated with antibiotics) yielded Bonferroni p-value = 2.5939e-12 and Holm p-value = 1.7293e-12; II vs III yielded Bonferroni p-value = 2.6645e-15 and Holm p-value = 2.2204e-13; II vs IV yielded Bonferroni p-value = 1.9363e-11 and Holm p-value = 9.6816e-12; III vs IV yielded Bonferroni p-value = 2.0811e-07 and Holm p-value = 6.9369e-08. When only pairs relative to I were simultaneously compared, Bonferroni and Holm results have shown that I vs II, I vs III, and I vs IV have yielded highly significant differences with varied Bonferroni p-value and Holm p-value.

## Discussion

In contrast to other types of injury, burn wounds stimulate metabolic and inflammatory adjustments that affect the patient to sundry impediments. Subsequent to burn injury, the innate immune system counters instantaneously by inducing localized and systemic inflammatory reactions. The innate immune response partakes in triggering the adaptive immune response; conversely, in doing so it has an undesirable consequence on the burn victim’s ability to build up a strong immune reaction to overrunning microorganism and, as a result, prompts the burn victim to infectious snags. Infection is the predominant reason of morbidity and death in burn-injured populations, with all most 61% of deaths being caused by infection [20]. Just after the burn incident, the wound surfaces are sterile but these wounds sooner or later become colonized with microorganisms. Bacteria like Gram-positive staphylococci, inhabiting sweat glands and hair follicles, can escape burn insult and eventually colonize the burn-affected area in absence of any topical treatment with antimicrobial agents. Sooner or later, other microbes including Gram-positive, Gram-negative, and yeasts colonizes these wounds [21]. *P. aeruginosa* is such an opportunistic pathogen which causes ruthless, severe and chronic nosocomial infections in burn patients. The majority of failures in burn treatments are linked to inapt primary antibiotic therapy. The heavily-burned patients always run risk of wound infection and incidence of infection has a direct correlation with the degree of burn injury. Undoubtedly, topical therapy with the existing antimicrobial agents has considerably decreased the occurrence of invasive Pseudomonas-burn-wound-sepsis, but none of the agents sterilize the burn wound. Hence, chances of the pathogens to get away from the confinement of the burn-wound and invade viable tissue increases [22].

Several studies were conducted using herbal drugs (represents a main part in all traditional systems of medicine) aiming prevention of infection and early wound healing. There are about 1250 Indian medicinal plants [23]; of which 46 plants (belonging to 44 genera and 26 families) were known to heal wounds and related injuries including burn wounds [24]. In one of the studies, burn injured rats treated with two Ayurvedic formulations, Jatyadighrita and Jatyadi tail, has revealed that the burn wound healing potential of Jatyadi tail was comparable to silver sulfadiazine [25]. Another Ayurvedic preparation, made with *Azadirachta indica A. Juss* leaves, resin of *Shorea robusta Gaerth. F*, and oil of *Linum vsitatissimum L.*, called ‘Atasthadi Lep’ when applied to burn wound of rat the % wound-heal recovery was significantly better than those wounds treated with silver sulfadiazine but complete eradication of infection was not attained [26]. The present study was planned to elucidate progress of healing *P. aeruginosa* (multiple-antibiotic-resistant strain HW01) – infected - burn-wound with the fermented polyherbal formulation (AP-01) compared to untreated animals with infected burn-wound and treated with antibiotics (piperacillin/ tazobactam) to which the strain was found sensitive (Table 1).

The choice and treatment dose of the fermented polyherbal formulation AP-01 over the burn surface was determined from the qualitative swarming-prevention test (Fig. 1A,1B) and results of the MIC assay (Table 2) respectively. It is evident from this study that the animals suffered burn-injury to an extent which confirmed susceptibility to infection with MAR *P. aeruginosa* strain *HW01* and enabled us to control burn-wound infection or to evaluate therapeutic efficacy of AP-01. There was significant difference in the rate of wound healing between untreated and *P. aeruginosa* infected burn wounds (both Bonferroni p-value and Holm p-value is 0.0000e+00). When we looked at % residual wound area (PRWA) on 14th day and then compared the averages of two separate groups: (i) treatment of *P. aeruginosa*-infected wound with antibiotic, and (ii) treatment of *P. aeruginosa*-infected wound with polyherbal formulation AP-01, and used T-test calculator for 2 independent means, the t-value is −21.01294 and the p-value is <0.00001. The result is significant at p < 0.01 or 0.05 implying that Gr-VI (treatment of the infected burn-wound with AP-01) was better than Gr-IV (antibiotic-treated infected burn-wound) supporting Fig. 2C & 2D. Again, Tukey HSD p-value for the pair, Gr-II (untreated burn infected with *P. aeruginosa HW01*) vs Gr-VI (burn infected with *P. aeruginosa HW01* and treated with AP01) was 0.0010053. The increased CRP level ebbed faster to the normal level with control of inflammation in the animals treated with polyherbal AP-01 (group V and group VI) within day 8 when serum CRP became undetectable while for other groups of animals it was detectable (Table 3). The pathogen-load of MAR *P. aeruginosa HW01* (qualitative assay) was zero in the infected burn-wounds on the 5th day following burn in case of AP-01 treated animals’ group no.VI, while detectable numbers of viable bacteria were still present over the infected burn wounds of either untreated group or the group treated with piperacillin/tazobactam (Table 4).

## Conclusion

A novel re-purposed Ayurveda-based polyherbal formulation as powerful target-blast-arsenal against notorious multiple-antibiotic-resistant (MAR) pathogens has been developed. Since several different active plant-based components were put together in a single formulation, no bacterium in a sizable population finds an escape route to avoid the deadly assault because mathematical probability to recruit multiple mutations simultaneously tends to zero. Hence, our approach is in stark contrast to the present scenario of the development of resistance within no time against a newly discovered antibiotic. Findings of our animal study revealed success against MAR *Pseudomonas aeruginosa* infection on rat burn wounds.

Therefore, the Ayurvedic fermented polyherbal formulation AP-01 may be used in future generation antimicrobial formulation in successful and efficient wound therapy.

## Authors’ contributions

SC: Formal analysis; methodology; investigation; writing original draft. SS: Formal analysis; investigation; image capture. RC: Conceptualization; funding acquisition, formal analysis; project administration; supervision; writing original draft; review and editing.

## Acknowledgements

Authors are thankful to Dr. Min Bahadur, Professor and Dr. Tilak Saha, Assistant Professor, Department of Zoology, University of North Bengal for their active co-operation in procurement of animals and necessary animal ethical permissions from Department of Zoology, University of North Bengal. Authors are grateful to Doctor Subrata Roy, B.A.M.S. of J.B.Roy State Ayurvedic Medical College and Hospital for identification of plant parts for preparing polyherbal formulation AP-01.

